# Analysis of the *Borreliaceae* Pangenome Reveals a Conserved Genomic Archictecture Across Phylogenetic Scales

**DOI:** 10.1101/2024.01.07.574540

**Authors:** Jacob E. Lemieux

**Affiliations:** Division of Infectious Diseases, Massachusetts General Hospital, Departments of Medicine and Microbiology, Harvard Medical School

## Abstract

The Family *Borreliaceae* contains arthropod-borne spirochetes that cause two widespread human diseases, Lyme disease (LD) and relapsing fever (RF). LD is a subacute, progressive illness with variable stage and tissue manifestations. RF is an acute febrile illness with prominent bacteremia that may recur and disseminate, particularly to the nervous system. Clinical heterogeneity is a hallmark of both diseases. While human clinical manifestations are influenced by a wide variety of factors, including immune status and host genetic susceptibility, there is evidence that *Borreliaceae* microbial factors influence the clinical manifestations of human disease caused by this Family of spirochetes. Despite these associations, the spirochete genes that influence the severity and manifestations of human disease are, for the most part, unknown. Recent work has identified lineage-specific expansions of lipoproteome-rich accessory genome elements in virulent clones of *B. burgdorferi*. Using publicly available genome assemblies, I show here that all *Borreliaceae* lineages for which sufficient sequence data is available harbor a similar pattern of strongly structured, lineage-specific expansions in their accessory genome, particularly among lipoproteins, and that this pattern holds across phylogenetic scales including genera, species, and genotypes. The relationships among pangenome elements suggest that infrequent episodes of marked genomic change followed by clonal expansion in geographically and enzootically structured populations may account for the unique lineage structure of *Borreliaceae*. This analysis informs future genotype-phenotype studies among *Borreliaceae* and lays a foundation for studies of individual gene function guided by phylogenetic patterns of conservation, diversification, gain, and/or loss.

## Introduction

Spirochetes of the Family *Borreliaceae* cause two major clinical syndromes: Relapsing fever (RF) and Lyme disease (LD). RF is an acute, febrile disease with prominent and often recurrent spirochetemia. Bloodstream survival of spirochetes, a hallmark of RF, is facilitated by sophisticated systems for antigenic variation [1]. LD, in contrast, is a subacute, progressive illness with variable, organ-specific manifestations, a consequence of the tropism of the spirochete to various host tissues [2]. LD and RF also have different primary tissue tropisms: RF spirochetes are found at the highest abundance in the bloodstream [3], whereas most LD spirochetes are found at the highest concentrations in tissues and occur only rarely, and at low levels, in the bloodstream [2].

The clinical divergence between LD and DRF is associated with a deep phylogenetic split between RF and LD spirochetes. More broadly, there is a general correlation between phylogenetic distance and clinical divergence among *Borreliaceae*: The largest phenotypic differences are observed between RF and LD spirochetes; lesser but still notable clinical variation is observed between species within a genus,^1^ and more subtle yet meaningful variation occurs at the genotype level. For example, all *Borreliella* spirochetes cause LD, but the clinical manifestations of LD vary based on the infecting genotype. *B. burgdorferi* is strongly associated with LA [9]; *B. garinii* and *B. bavariensis* are strongly associated with LNB [10–12], and *B. afzelii* is associated with ACA [10–12] and *Borrelial* lymphocytoma [13,14]. At the genotype level, which is best studied for *B. burgdorferi* sensu stricto, OspC type A / RST 1 genotypes of *B. burgdorferi* disseminate in humans at higher rates [15–18], cause higher rates of symptoms [19], greater inflammation [19,20], and more severe LA [19]. Thus, the divergence of clinical disease in humans correlates with genetic divergence across phylogenetic scales in *Borreliaceae*.

Despite strong evidence that microbial genetic factors influence clinical manifestations, the specific genes that contribute to these phenotypic differences are incompletely understood. Many genes lack annotated functions; in other caess, where some or all functional activities are well-characterized, including systems for antigenic variation, serum resistance and and tissue adhesion (reviewed in [21–23]), gaps remain in understanding how these functions work together to influence disease manifestations. This is in part due to the complexity of the *Borreliaceae* genomes, which contain numerous plasmids that are challenging to sequence and annotate [24,25], and the relative rarity of human clinical isolates of LD and RF, which require specialized culture conditions not typically performed in routine clinical practice [26].

Identifying genes that contribute to distinct clinical manifestations of LD and RF remains an important scientific goal because of its potential to reveal fundamental insight into *Borreliaceae* pathogenesis and open new opportunities for therapy and prevention. Recent work has used microbial genome-wide association methods to begin to identify the genetic elements associated with clinical manifestations of LD [18] but have not yet resolved individual causal genes from among sets of linked loci. Phylogenetic patterns across *Borreliaceae* can help prioritize individual genes for functional follow-up, and extending genotype-phenotype studies to multiple *Borreliaceae* linages requires an understanding of genomic relationships and extant genomic data for the Family. Here, I use publicly available genome sequence data to construct pangenomes and characterize lineage-specific genomic events from across *Borreliaceae*.

## Materials and Methods

*Borreliaceae* genome assemblies and associated metadata were downloaded from NCBI. Assemblies were filtered to be be > 1 Mb (excluding chromosome-only assemblies) and < 2 Mb. Assemblies were annotated using Prokka [27]. Pangenomes were constructed using Roary [28], with a minimum BLAST threshold of 80% homology and without splitting of paralogous groups, and PIRATE [29] using default parameters. Trees were constructed using FastTree [30] and IQ-TREE2 [31] with 1000 ultrafast bootstrap replicates. The pipeline was executed using Snakemake [32]. Lipoproteins were annotated using SpLip [33]. The resulting pangenomes, phylogenetic trees, and metadata files were analyzed and plotted using R [34], ggplot [35] and ggtree [36]. The genomic position of pangenome homology groups was annotated by aligning pangenome elements against *Borreliaceae* reference genomes using minimap2 [37]. Because chromosome sequences are easier to assemble than plasmid sequences in *Borreliaceae* [24,38], and therefore presumably more accurate if assemblies are unfished, I used the following approach: If pangenome elements aligned to the chromosome sequences, they were annotated as provisionally chromosomal. If they aligned to reference sequences but not to chromosome sequences, they were annotated as provisionally plasmid-encoded. Lineage-specific associations of individual pangenome elements were identified using logistic regression to predict the lineage based on the pangenome homology group, as implemented in R using the glm() command.

Ethical approval: The analysis of microbial genome sequences from deidentified patient data was approved by the MGH Institutional Review Board under protocol 2019P001846.

## Results

To characterize the pangenome relationships among *Borreliaceae,* I downloaded all publicly available *Borreliaceae* assemblies from NCBI Genbank, annotated their genomes, and constructed a pangenome and phylogenetic tree from core genome sequences. As of December 26, 2023, NCBI Genbank contained 702 *Borreliaceae* genome assemblies from 20 *Borreliella* taxa and 18 *Borrelia* taxa from both the RF group and the “reptilian” (REP) group [39]. The core genome tree constructed from these assemblies is shown in Figure 1, with overrepresented taxa downsampled to a maximum of 25. Consistent with prior phylogenetic analyses, the tree reveals a well-supported, deep division into two monophyletic branches consisting of *Borrelia spp.* and *Borreliella spp.* spirochetes. *Borrelia spp.* contain two monophyletic sister clades, RF spirochetes, and the REP group.

**Figure 1:**
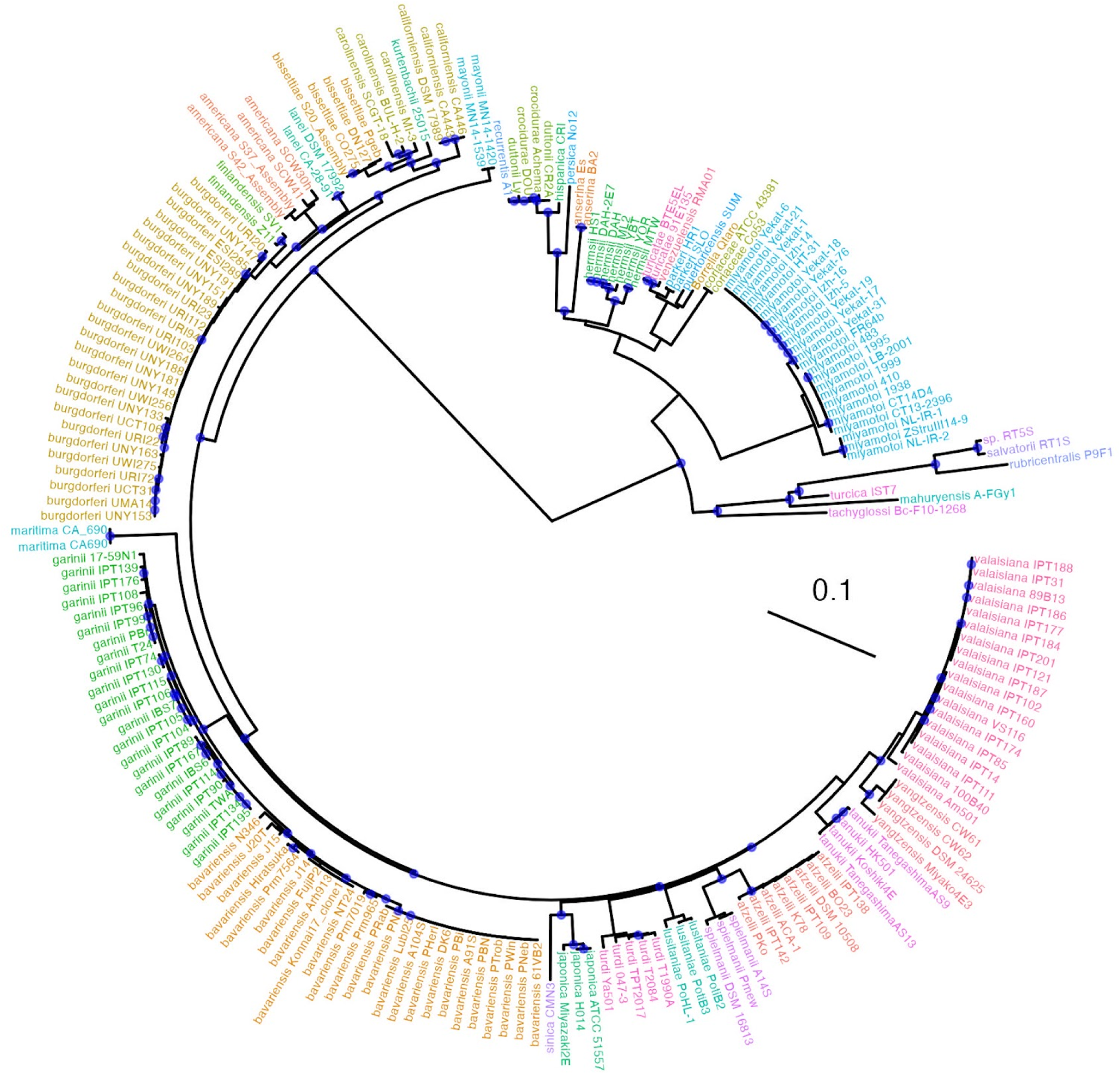
Core Genome Phylogenetic Relationships Among Borreliaceae. Radial tree of *Borreliaceae* with tips colored by species. Nodes with bootstrap support > 90% are colored with a blue dot. The scale bar denotes nucleotide substitutions per site.

The *Borreliaceae* pangenome contains 299 core ORFs at a protein BLAST homology threshold of 80% (Figure 2). The non-core (accessory) pangenome among *Borreliaceae* contains 5684 ORF homology groups, although this estimate is probably inflated due to fragmented assemblies [40]. The accessory genome is strongly structured by lineage, with divisions in the accessory genome following the core genome phylogeny. The deep phylogenetic division between *Borrelia spp.* and *Borreliella spp*. is mirrored in the divergence of the accessory genomes. *Borrelia spp.* and *Borreliella spp.* possess additional genus- and species-specific core genomes that are readily apparent in Figure 2. Annotated pangenomes can be used to define genus- and species-specific pangenomes and lipoproteomes (Supplemental File 1). This analysis was also repeated using PIRATE [29], which allows for the construction of pangenomes across multiple homology, confirming that this accessory genome structure was not an artifact of a single homology threshold.

**Figure 2:**
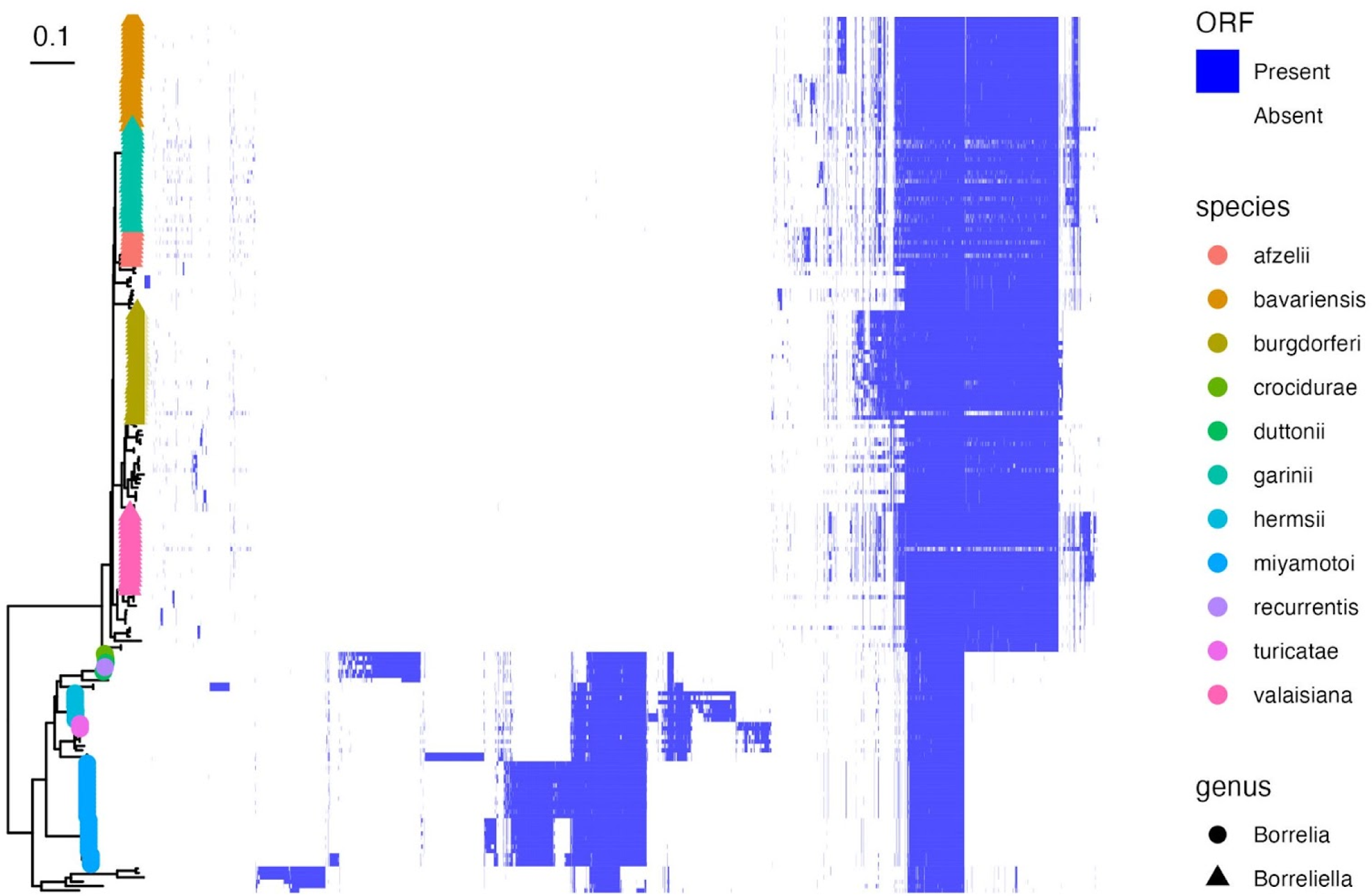
Pangenome elements by core genome phylogeny. A matrix showing the presence or absence of pangenome homology groups in each strain is shown. Pangenome homology groups that are present are denoted in blue. Groups that are absent are uncolored. Each row corresponds to an individual strain and each column corresponds to individual homology groups. The rows are ordered according to a phylogenetic tree, with tip shapes and colors labeled by the *Borreliaceae* genus and species. The columns are clustered with hierarchical clustering. The tree scale bar denotes nucleotide substitutions per site.

The structure of accessory genome elements consists of a lineage-specific “block” pattern in which groupings of accessory genome presence are almost perfectly correlated with lineage-defining subdivisions in the core genome tree (Figure 2). This pattern closely resembles the pattern observed for the *B. burgdorferi* sensu stricto pangenome [18] but is present across phylogenetic scales. Within the *Borreliella spp.,* there are lineage-specific groups of accessory loci, for example among *B. burgdorferi* (*B. burgdorferi* sensu stricto), *B. garinii*, *B. bavariensis*, and *B. afzelii* genomes. *Borrelia spp.* similarly possess lineage-specific changes such that there are tightly-structured core genomes for Old World RF agents (including *B. crocidurae* and *B. duttonii*), hard-tick RF agents (*B. miyamotoi*), and New World RF agents (including *B. turicatae, B. parkeri,* and *B. hermsii*). Within sublineages, a similar pattern occurs. For example, *B. miyamotoi* also shows evidence of distinct genotypes, with subsets of *B. miyamotoi* diverging based on geographic origin [41]. Russian and Asian *B. miyamotoi* isolates possess a slightly enlarged pangenome and a distinct block of genetic elements compared to those from Western Europe and North America, each of which forms a distinct clade (Figure S1). Among *B. burgdorferi* sublineages, OspC Type A / RST1 genotypes possess a lineage-core group of ORFs, as shown in reference [18].

Lipoproteins are particularly important for *Borreliaceae* immune evasion, tissue tropism, and other aspects of vertebrate pathogenesis [2,42]. To characterize the relationships among the *Borreliaceae* pan-lipoproteome, I annotated lipoproteins using SpLip [33] and analyzed the phylogenetic patterns of lipoprotein-annotated homology groups (Figure 3). The strong lineage-specific pattern of the pangenome (Figure 2) is reproduced almost exactly in the lipoproteome (Figure 3), except that all lipoproteins are lineage-variable at the homology thresholds used.

**Figure 3:**
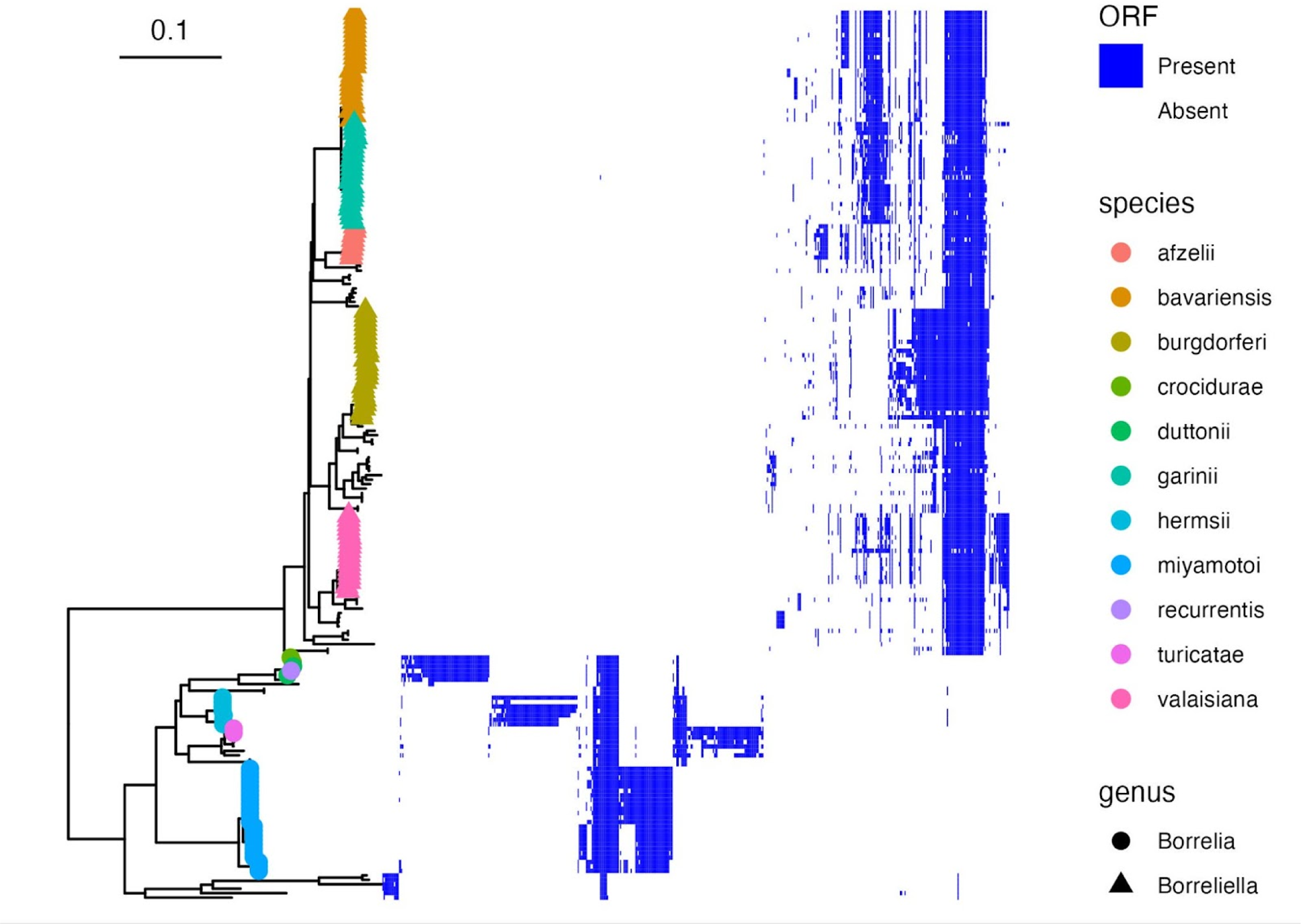
The *Borreliaceae* lipoproteome. A matrix showing the presence or absence of probable lipoproteins, as classified by SpLip [33], in each strain is shown. Each row corresponds to an individual strain and each column corresponds to a homology group classified as a probable lipoprotein. Rows are ordered by core genome phylogeny and columns are ordered by hierarchical clustering.

I next assessed the spatial organization of homology groups within *Borreliaceae* genomes. Because many assemblies are incomplete due to the difficulty of generating complete assemblies through short-read sequencing [24,25,43], I conducted a provisional analysis of draft genome organization by aligning a representative of each pangenome homology group to well-annotated reference genomes. Lipoproteins are preferentially encoded on plasmids throughout *Borreliaceae* (Figure 4). Core genome elements were almost exclusively encoded on the chromosome (Figure S2). Accessory genome elements are preferentially encoded by plasmids (Figure S3), including all *Borreliella* accessory genome elements and many *Borrelia* accessory elements. One notable difference between *Borrelia spp.* and *Borreliella spp.* is that in *Borrelia spp.* portions of the species-specific core genome are encoded on the chromosome (Figure S2), whereas in *Borreliella* all nearly all species-specific core genomes are chromosomal (Figure S3). Thus, the pattern of lineages defined by plasmid-encoded accessory elements enriched in lipoproteins is present but less absolute in *Borrelia* compared to *Borreliella*. Together, these analyses enable the identification of lineage-specific lipoproteomes (Figure S4-S5), a resolution which can highlight more subtle lineage-specific genomic patterns such as the unique sequences associated with OspC Type A / RST1 genotypes among *B. burgdorferi* (Figure S4B-C) [18].

**Figure 4:**
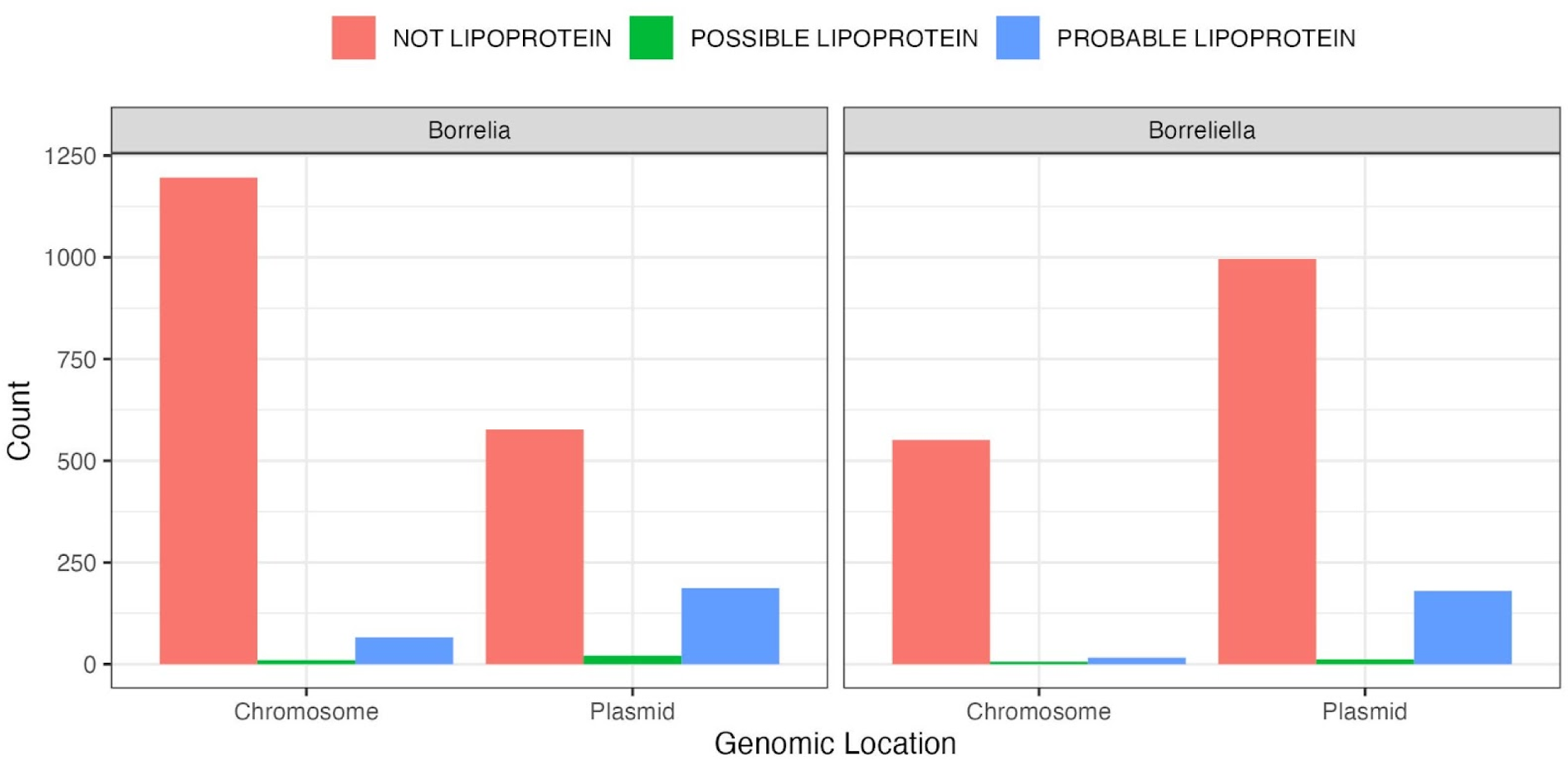
Genomic location of pangenome elements. The genomic location of pangenome elements is shown. Counts are stratified by whether they are classified by SpLip [33] as probable lipoproteins, possible lipoproteins, or not lipoproteins.

By summing the number homology groups in an individual genome, pangenome analysis also provides an estimate of the size of the coding genome and lipoproteome. *Borrelia* genomes have slightly more ORF homology groups than *Borreliella* genomes (Figure 5A), an effect which is driven by soft tick *Borrelia* RF spirochetes (Figure 5B), particularly among lipoproteins (Figure 5C-D). Differences are seen among arthropod vectors. The hard tick RF spirochete *B. miyamotoi* has fewer ORF homology groups and probable lipoproteins than soft tick RF spirochetes. Despite being an RF spirochete, its number of ORFs and probable lipoproteins are similar to LD spirochetes. Among LD *Borreliella, B. garinii* and *B. bavariensis* have fewer ORF homology groups than *B. burgdorferi;* however, there is marked variation in genome size within *B. burgdorferi* (Figure 5 and [18]) and other species (Figure 5B, 5D).

**Figure 5:**
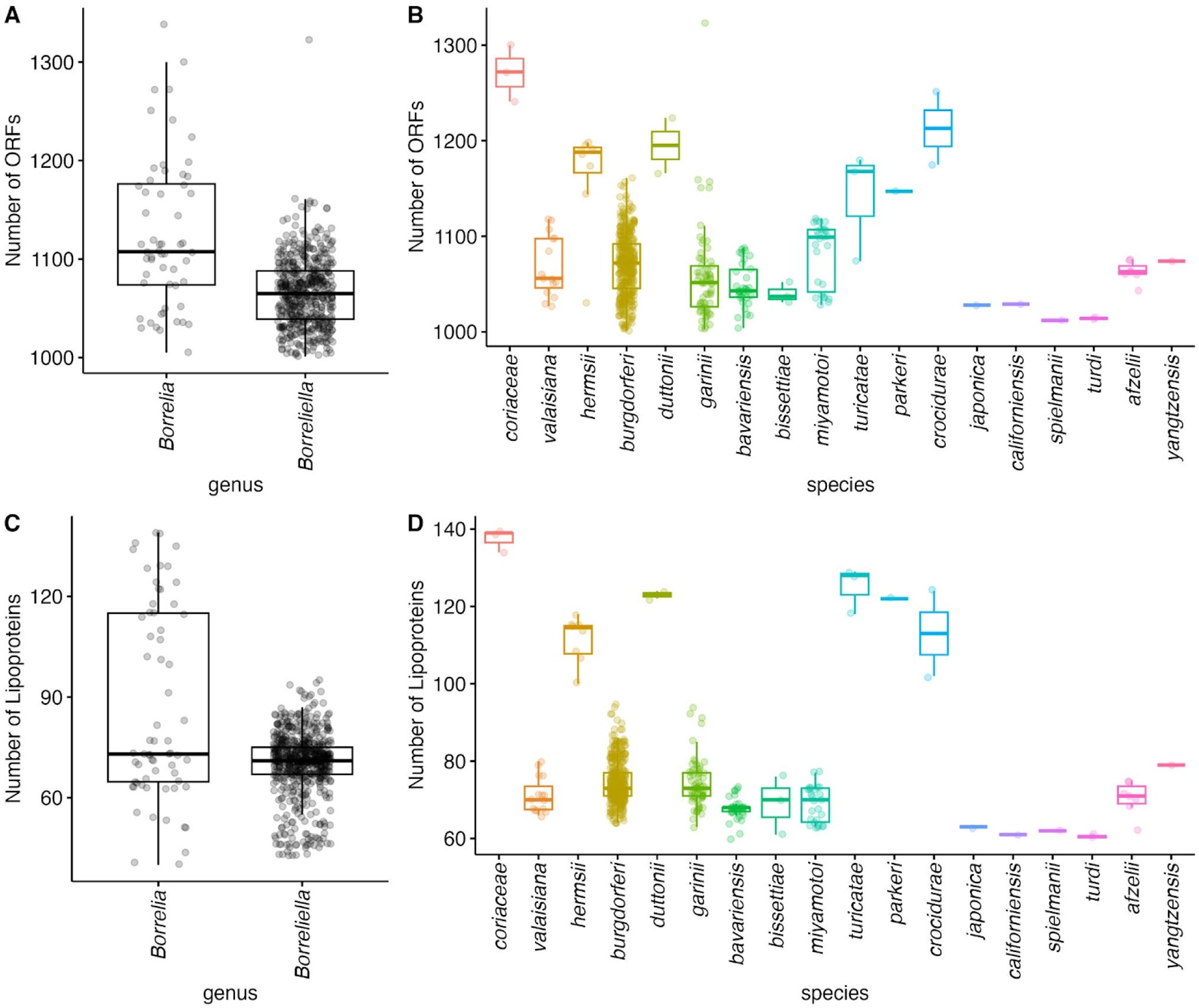
Coding genome and lipoproteome size by genus and species. **A.** The number of ORF homology groups for each genome assessed, grouped by genus. **B.** The number of ORF homology groups for each genome assessed, grouped by species. **C.** The number of ORF homology groups annotated as probable lipoproteins for each genome assessed, groupbed by genus. **D.** The number of ORF groups annotated as probable lipoproteins for each genome assessed, grouped by species.

## Discussion

The growing quantity of genomic data in publicly available data banks provides unique opportunities to characterize the evolutionary genomic relationships among *Borreliaceae*. Such characterizations can help prioritize individual genes for further investigation and provide a basis for genome-wide studies of genotype-phenotype correlations. Recent work has shown that lineages of *B. burgdorferi* contain groups of strongly linked loci accessory elements, particularly surface lipoproteins encoded on plasmids, and these blocks of loci are linked to clinical manifestations [18]. Using published assemblies of *Borreliaceae*, I show here that this pattern extends across phylogenetic scales and appears to be a general organizing principle of the *Borreliaceae* pangenome. The large number of genomes included from across the Family is a strength of this analysis and builds on previous reports [18,44], providing a panoramic view of *Borreliaceae* genome diversity. Limitations include a convenience sample with a lack of standardization among sequencing, assembly, and submission procedures; incomplete and fragmented assemblies that may inflate the number of pangenome elements [40] and do not easily permit an analysis of associated genomic context; and a focus on variation at the level of homology clusters rather than single genetic variants.

The strong block structure of accessory genome elements across phylogenetic scales is remarkable. The monophyletic nature of these blocks implies that most have a single common ancestor and that the set of genes included in the block has been stable over evolutionary time. This pattern suggests that extant variation in *Borreliaceae* is the consequence of a relatively small number of profound ancestral changes involving multiple genetic elements, and that these marked changes occurred simultaneously or in short succession.

The abrupt genesis of major genomic changes is consistent with a “punctuated equilibrium” model[45] of evolution. This pattern would the strong statistical linkage between physically unlinked markers, e.g. OspC and the *B. burgdorferi* ribosomal spacer [46].

It is notable that the patterns holds for major and minor lineages across scales. The fractal nature of this pattern may provide clues into the underlying mechanisms that result in this pattern. Fractals occur when the same laws apply recursively across scales [47]. The process of lineage generation may be an example of such a recursive law and suggests the following mechanism: the creation of a new genotype begins with a rare genomic event that involves the gain, loss, and/or exchange of multiple genes, probably in an arthropod or vertebrate host infected with multiple genotypes and/or different species. Such a lineage may clonally expand in a new niche, leading to geographically and/or ecologically structured populations. Over time, these populations accumulate further genetic diversity, eventually become differentiated enough to be classified as a species or genus. Along the way, sublineages form recursively by the same mechanism.

Although this mechanism raises questions about how *Borreliaceae* could survive abrupt genetic shifts, complex genomic rearrangements are increasingly recognized in other contexts, especially in cancer, where the phenonomen is termed chromothripsis [48–50]. The unique genome organization of *Borreliaceae* [24,51]—in which conserved functions are encoded by a single chromosome and genes that mediate host-pathogen interactions, particularly surface lipoproteins are encoded on plasmids—probably facilitates organism survival following an episode of dramatic change by allowing loss, gain, or exchange of multiple genes on plasmids while insulating the core metabolic or housekeeping functions of the spirochete on the chromosome and certain conserved plasmids. Other mechanisms may also predispose to complex genetic rearrangements: Both RF and LD *Borreliaceae* genomes contain sophisticated machinery for inducing gene conversation to support antigenic variation [21,52–54]. Regulation of this process is obviously paramount to genome integrity. Genomic instability involving multiple linked loci is readily observable in *B. burgdorferi*, in which routine *in vitro* culture is associated with spontaneous plasmid loss [55,56], and double-stranded breaks appear to result in plasmid loss *in vitro* [57].

The genetic structure of *Borreliaceae* informs the search for microbial loci associated with human disease in several ways. Lineage-level associations aggregate multiple genes’ worth of effects, potentially increasing the effect size of the genotype-phenotype link, and the consistent genetic profile among strains within a given lineage and reduce the variance of these effects because strains are nearly isogenic. Thus, lineage-level associations should be relatively easy to identify, which may be why so many are known. On the other hand, locus-level effects will be more difficult to identify because individual loci are rarely observed alone. Large samples will be required, and statistical techniques, such conditioning on lineage-level effects, may help. Experimental methods in appropriate disease models involving reverse genetics will likely be necessary to assign or exclude causal roles for individual genes. Functional genomic approaches capable of screening multiple loci, such as TnSeq [58], CRISPRi using dCas9 [59,60], and gain-of-function screens [61,62] have recently been developed and are well-suited to the task of identifying experimentally important loci from among a larger set of candidates. Lineage-level associations greatly narrow the list of candidates for functional characterization and reverse genetic methods.

In summary, analysis of *Borreliaceae* pangenomes reveals a conserved genomic architecture in which lineages are defined by correlated blocks of accessory genome elements, primarily encoded on plasmids and enriched for lipoproteins, a pattern which holds across phylogenetic scales. This unusual fractal genetic structure provides clues into the origin and evolution of *Borreliaceae* lineages and establishes a basis for reductionist studies of gene function informed by Family-wide phylogenetic relationships.

## Acknowledgments

I gratefully acknowledge all the sequence submitters and contributors who deposited *Borreliaceae* assemblies in Genbank (Supplemental File 2), as well as Klemen Strle, Ira Schwartz, and Allen Steere for helpful discussions. This work was supported by NIH award K99/R00AI148604.

## Code availability

Analysis code is available at github.org/jacoblemieux/panborreliaceae.

## Footnotes

1 Recent phylogenetic analysis supports separation of RF and LD agents into separate genera[4]. The authors of reference[4] proposed revising the taxonomic assignments, designating *B. burgdorferi* sensu lato as *Borreliella* while retaining *Borrelia* for RF spirochetes based on historical priority.[4] Although the proposed taxonomic revision is controverial[5–7], I follow NCBI here and use *Borrelia* to refer to RF spirochetes and *Borreliella* to refer to LD spirochetes, although use of these terms by Genbank has also been criticized [8]. My goal here is clarity and consistency with the data source, rather than to take a position on the nomenclature.

## SUPPLEMENTAL FIGURES

**Figure S1:**
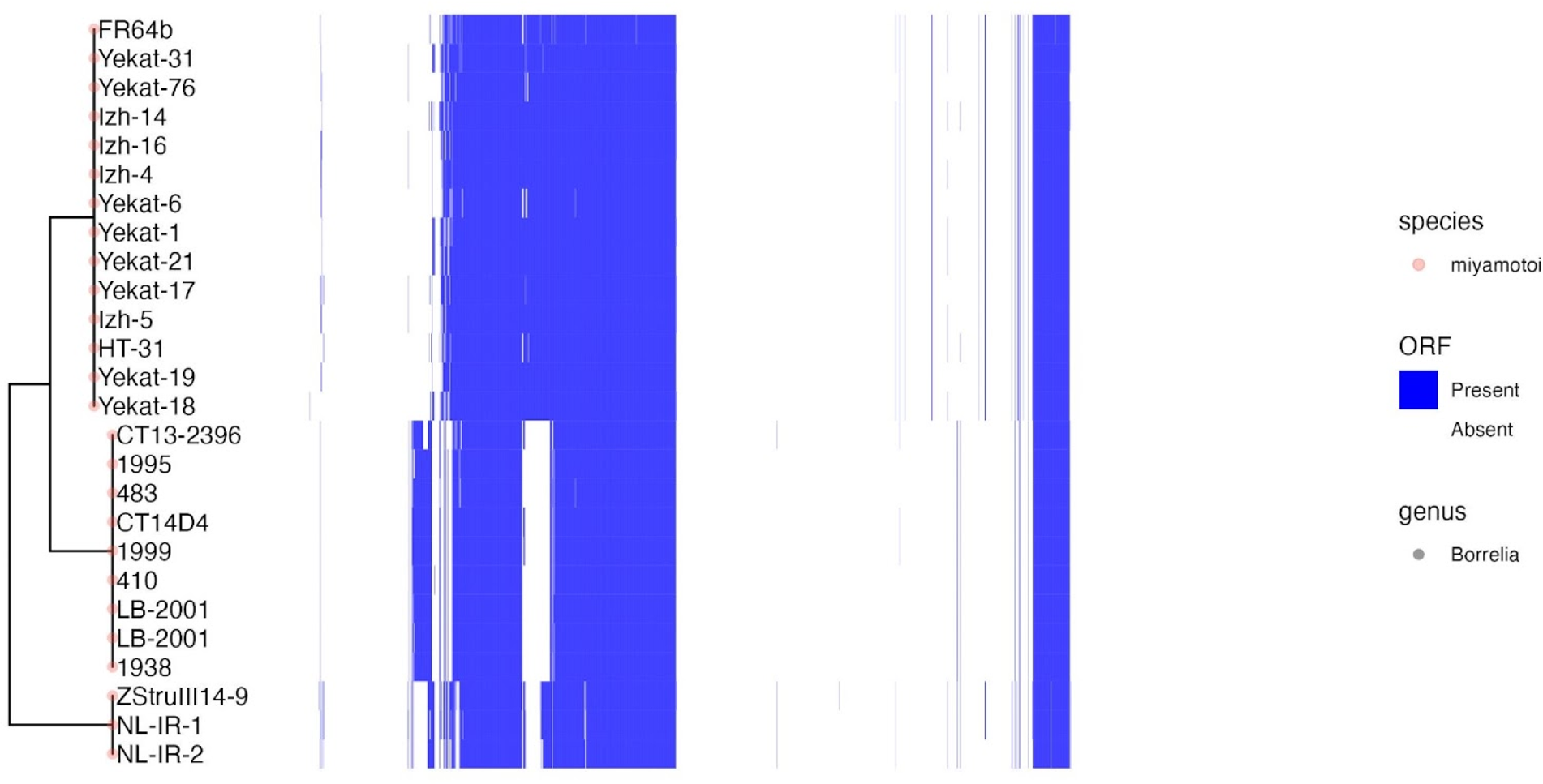
*B. miyamotoi* pangenome sequences showing the divergence of genotypes. The upper clade is composed of Russian and Asian strains. The middle clade is composed of North American strains. The bottom clade contains Western European samples. Pangenome homology groups that are present in a given strain are denoted in blue. Groups that are absent are uncolored. Each row corresponds to an individual strain and each column corresponds to individual homology groups. The rows are ordered according to a phylogenetic tree. The columns are clustered with hierarchical clustering.

**Figure S2:**
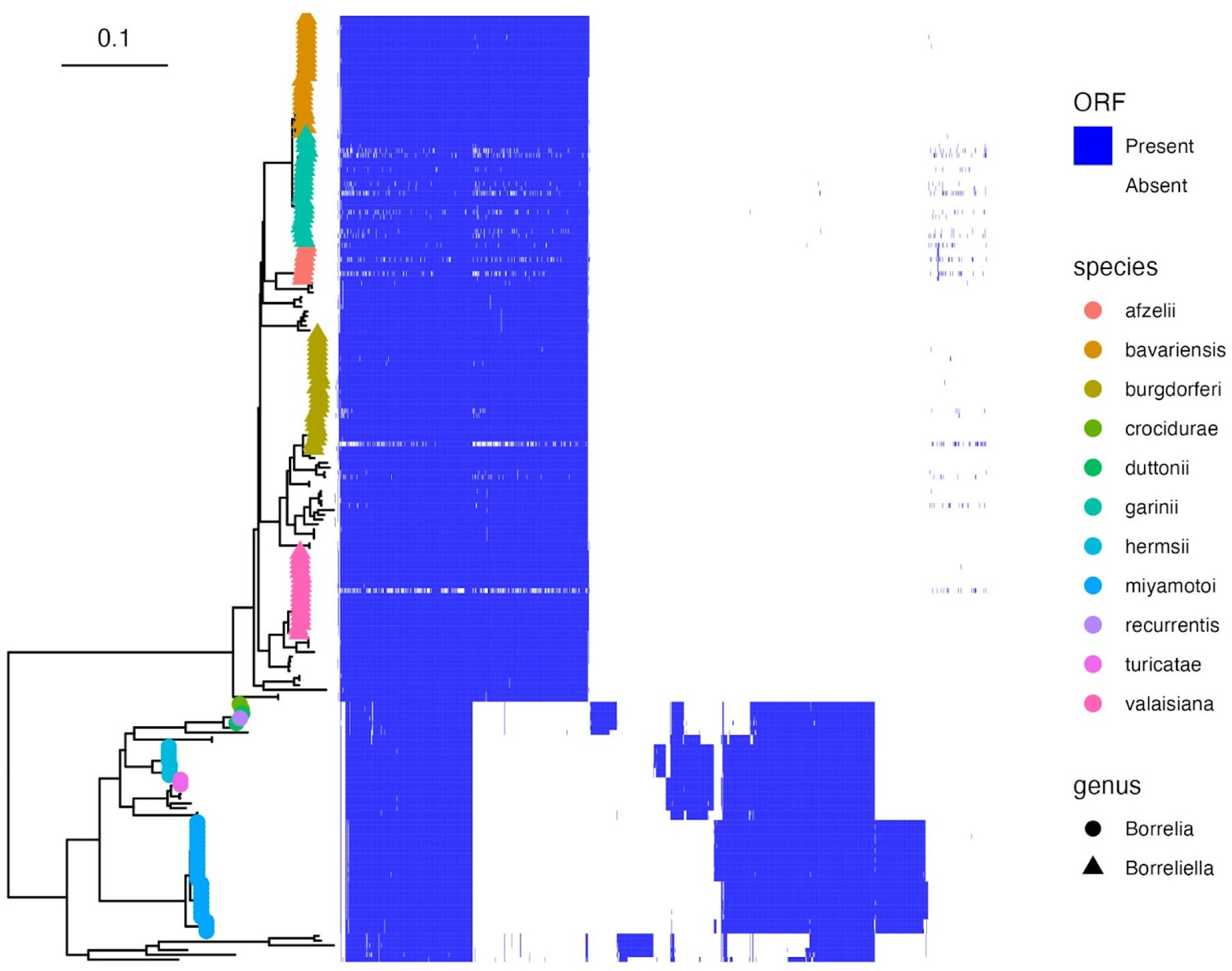
Chromosomally-encoded pangenome elements. Pangenome homology groups annotated as encoded on the chromosome are shown. Pangenome homology groups that are present in a given strain are denoted in blue. Groups that are absent are uncolored. Each row corresponds to an individual strain and each column corresponds to individual homology groups. The rows are ordered according to a phylogenetic tree, with tip shapes and colors labeled by the *Borreliaceae* genus and species. The columns are clustered with hierarchical clustering.

**Figure S3:**
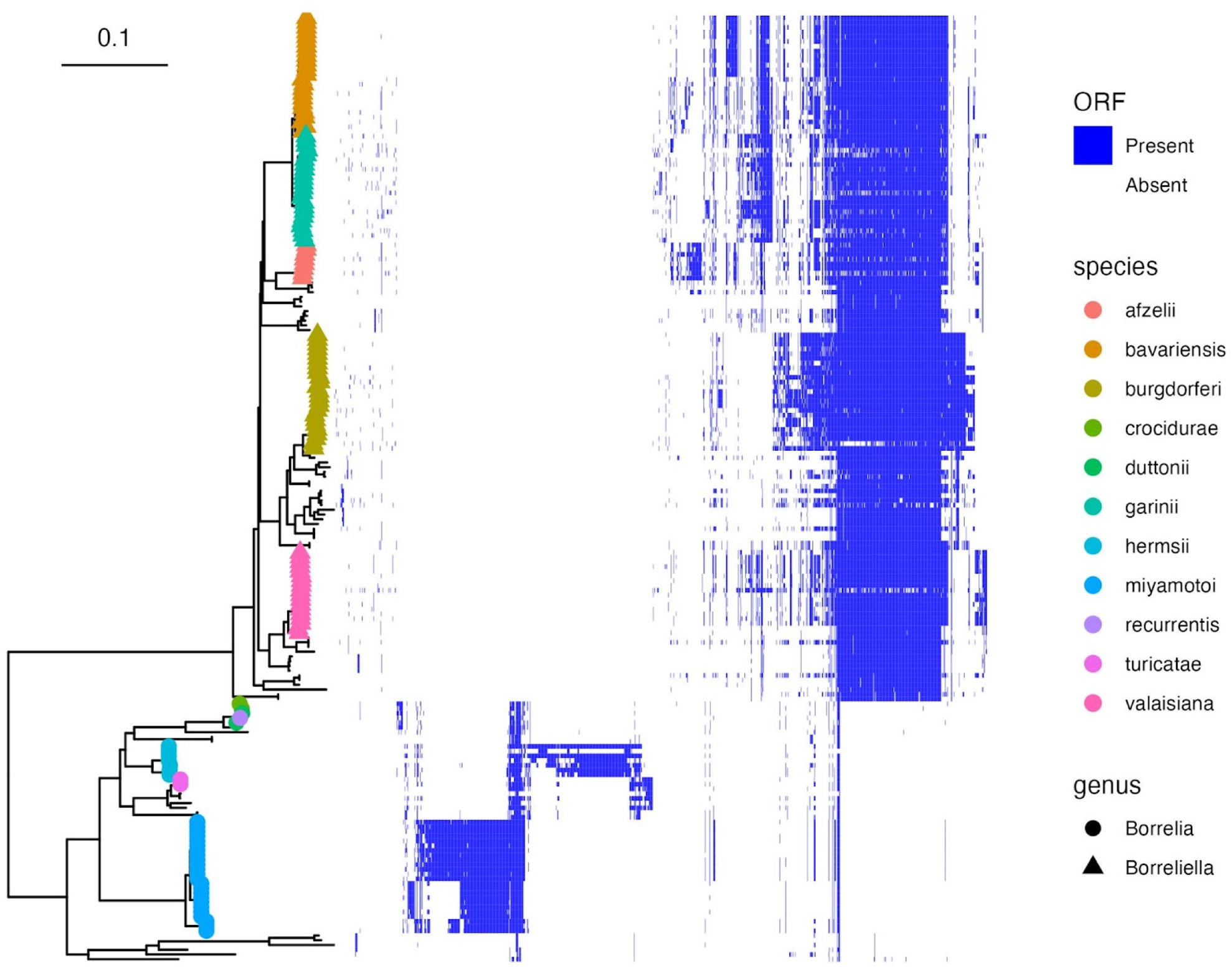
Plasmid-encoded elements. Pangenome homology groups annotated as encoded on plasmids are shown. Homology groups that are present in a given strain are denoted in blue. Groups that are absent are uncolored. Each row corresponds to an individual strain and each column corresponds to individual homology groups. The rows are ordered according to a phylogenetic tree, with tip shapes and colors labeled by the *Borreliaceae* genus and species. The columns are clustered with hierarchical clustering.

**Figure S4:**
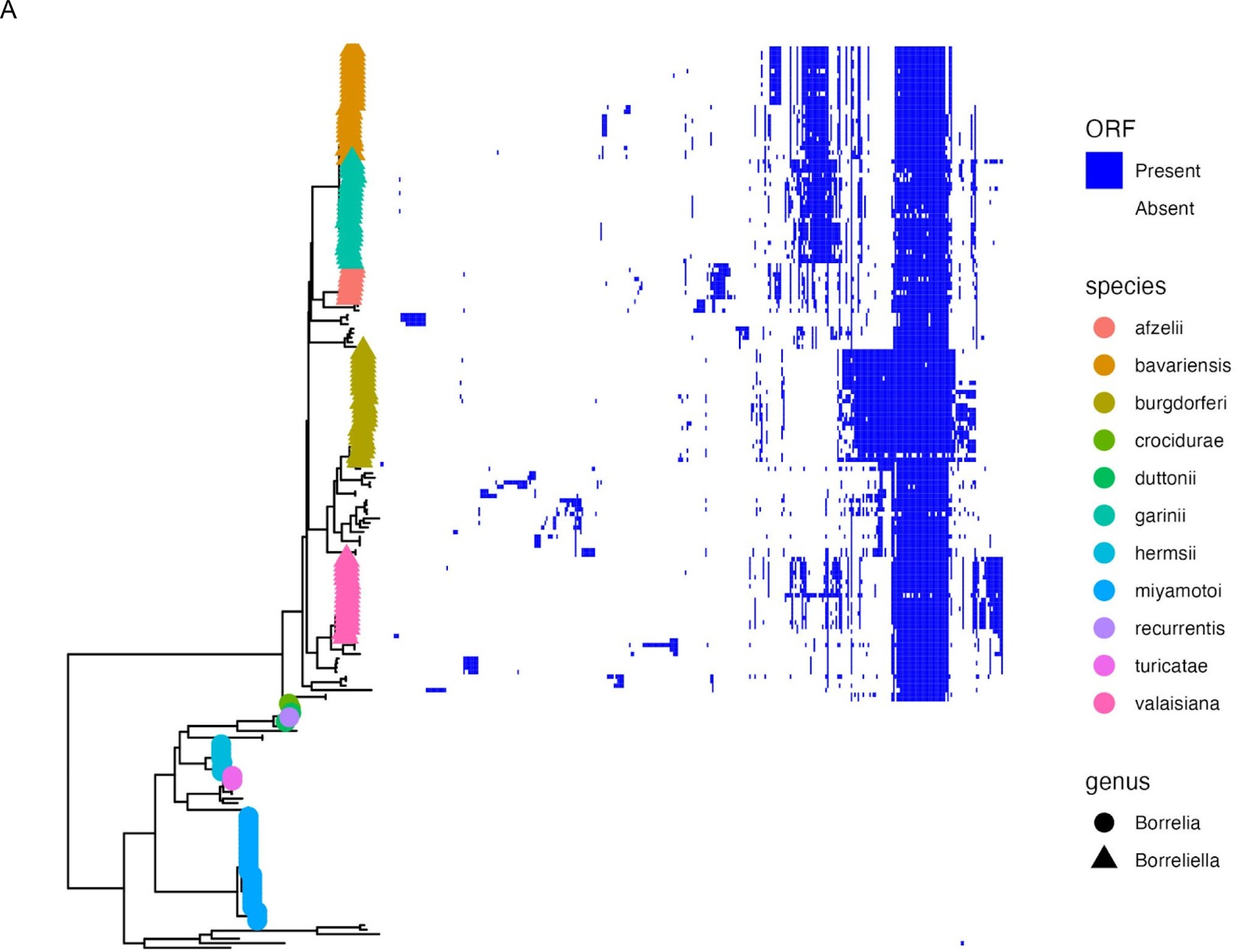
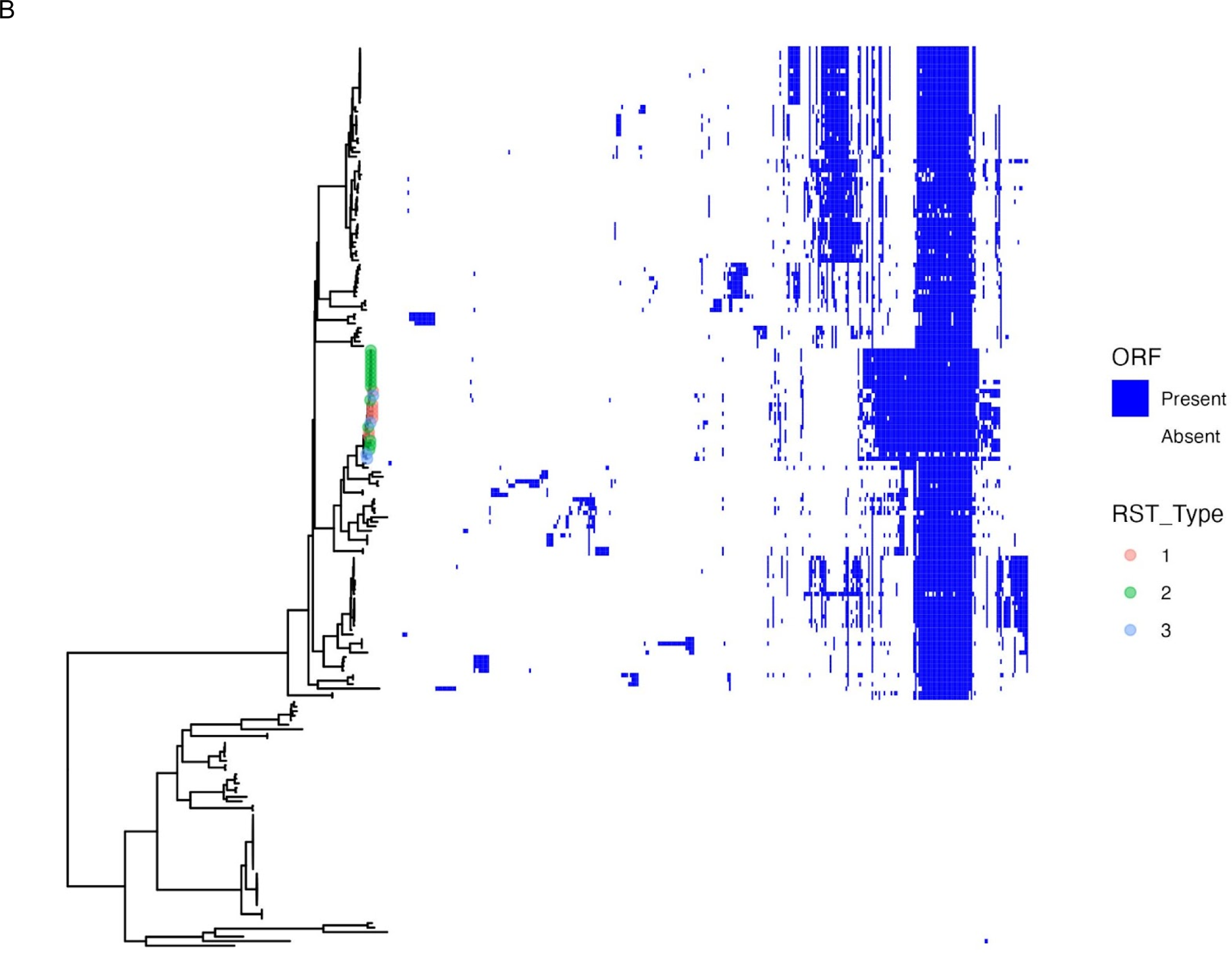
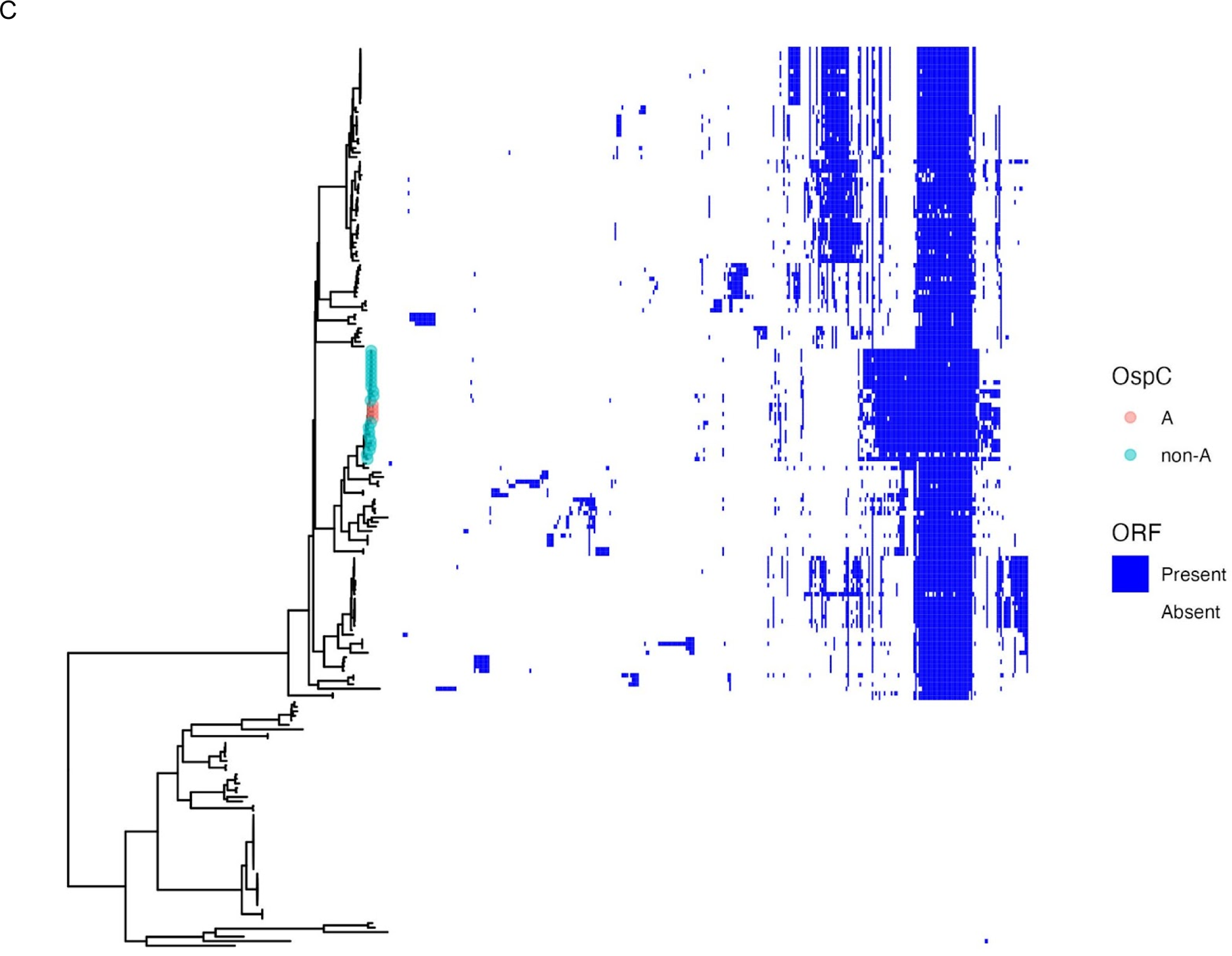
Phylogenetic and pangenome relationships of *Borreliella-*associated lipoproteins, by species and genotype markers. **A.** Core genome phylogenetic tree and presence/absence matrix of *Borreliella*-associated lipoproteins. Pangenome homology groups that are present are denoted in blue. Groups that are absent are uncolored. Each row corresponds to an individual strain and each column corresponds to individual homology groups. The rows are ordered according to a phylogenetic tree, with tip shapes and colors labeled by the *Borreliaceae* genus and species. The columns are clustered with hierarchical clustering. **B.** Tree and presence/absence matrix as in panel A with *B. burgdorferi* tips labeled according to RST type. **C.** Tree and presence/absence matrix as in panel A with *B. burgdorferi* tips labeled according to whether OspC type is A or not.

**Figure S5:**
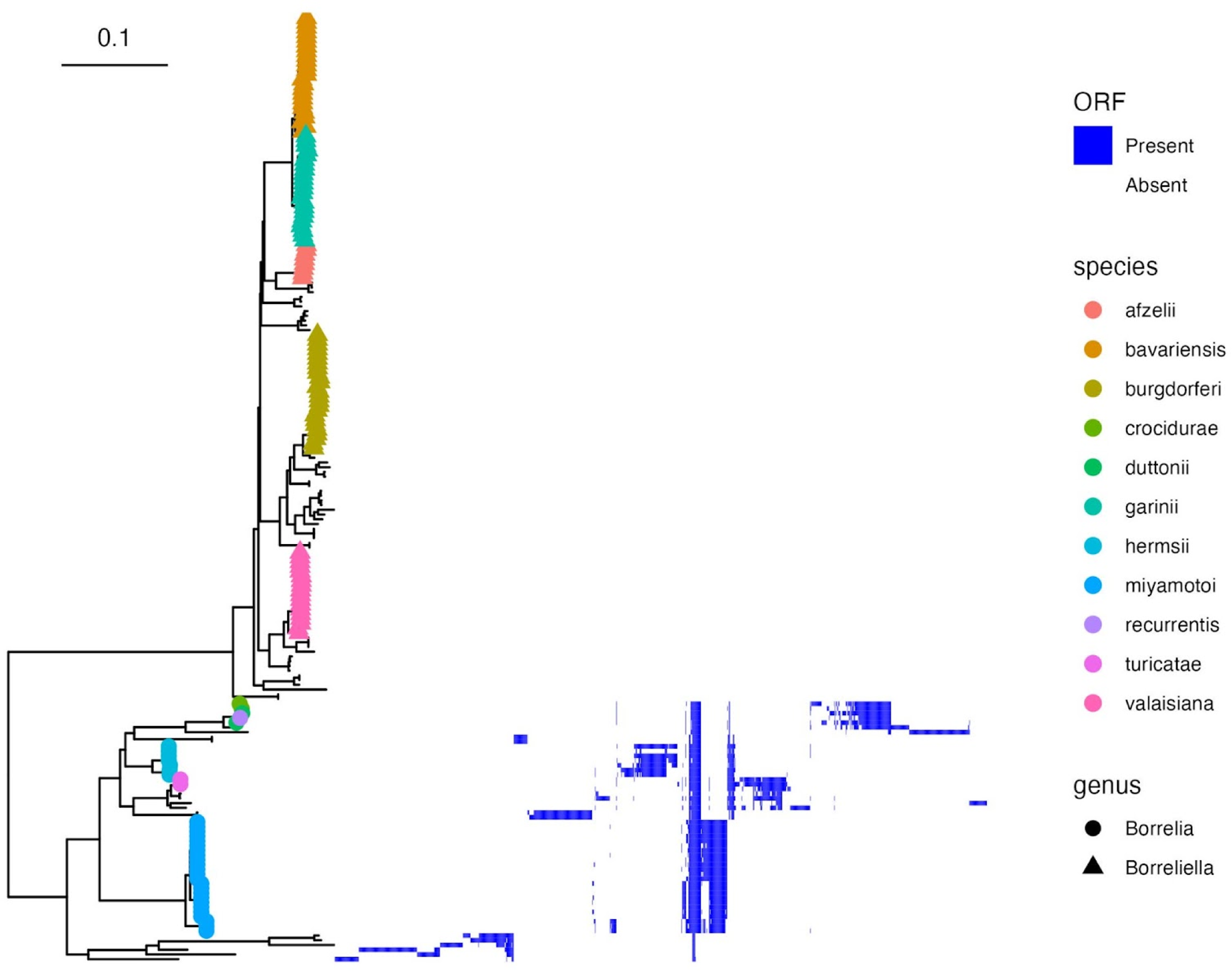
*Borrellia*-associated lipoproteins. Pangenome homology groups that are present in a given strain are denoted in blue. Groups that are absent are uncolored. Each row corresponds to an individual strain and each column corresponds to individual homology groups. The rows are ordered according to a phylogenetic tree, with tip shapes and colors labeled by the *Borreliaceae* genus and species. The columns are clustered with hierarchical clustering.

